# Robust mammalian RNA localization elements are complex and multipartite

**DOI:** 10.64898/2026.06.09.731215

**Authors:** Charlie Moffatt, Sharifah Syed Salim, Karl Bauer, Andrea MacFadden, Alli Jimenez, Chase Weidmann, Allison W McClure, Daniel Dominguez, Michael Kiebler, J. Matthew Taliaferro

## Abstract

The subcellular localization patterns of RNAs are controlled by regulatory elements contained within them. However, for most localized RNAs, the identities of these elements remain unknown. We had previously identified several localization elements that are necessary and sufficient for robust, kinesin-dependent RNA targeting to microtubule plus ends in a variety of cell types. Yet the characteristics of these elements that are critical for function remained unclear. To address this, we systematically created tens of thousands of mutant localization elements and quantified their ability to regulate subcellular RNA localization in neuronal cells. We found that the minimally active size of these localization elements is large, approximately 200 nucleotides. These elements contain multiple important subsequences, with some being completely intolerant of any changes and others being tolerant to a shuffling of nucleotide order but not to changes in nucleotide composition. Using single molecule microscopy, we verified these findings in primary rat neurons. Together, these results demonstrate that highly active mammalian RNA localization elements are large, complex, and multipartite and lay a foundation for further mechanistic studies of their function.

## INTRODUCTION

The subcellular localization of specific RNA molecules is critical for proper developmental patterning [1–6] and neuronal function [7–14]. Although hundreds of RNAs are subcellularly localized, the mechanisms underlying this localization for the vast majority of localized RNAs are unknown [15]. The handful of well characterized examples of RNA localization form the basis of the most common model of RNA localization, where *cis*-elements within trafficked RNAs are recognized by *trans*-factors that interact with the cellular transport machinery [16].

For most localized RNAs, the *cis*-elements, also called localization elements (LEs) or “zipcodes,” that regulate their transport are unknown. Of the relatively few well characterized LEs, many of them are large in size. For example, the LEs within the *D. melanogaster* K10, *oskar*, and *bicoid* RNAs are 44 nucleotides (nt), 32 nt, and 625 nt long, respectively [2,6,17–20]. This stands in contrast to elements that regulate other RNA metabolic processes like pre-mRNA splicing or translation which are typically between 4 and 12 nt long [21,22].

For many known LEs, their RNA structures are critically linked to their ability to regulate RNA transport. Again, the best characterized examples come from *Drosophila*. The K10 transcript localization signal (TLS) requires the formation of a 44 nt stem loop for localization activity [17,23]. The formation of a stem-loop structure after pre-mRNA splicing is required for the transport of *oskar* RNA [19,20]. The *bicoid* LE forms extensive secondary structure that is necessary for its interaction with Staufen and subsequent transport [6,24]. More recent work has also implicated the tertiary RNA structure of LEs in their ability to bind the protein cofactors necessary for their transport [25,26].

Currently known LEs were often identified by serially testing the localization of single reporters containing truncations or large deletions of a known active sequence. The localization of these reporters was typically assayed using imaging approaches. In some cases, the smallest active sequence identified was then subjected to mutational analysis [1–6]. The laborious nature of these experiments often limits the resolution of LE identity as each individual sequence tested requires its own round of design, cloning, and assaying.

With developments in high-throughput sequencing and large-scale *de novo* DNA synthesis, new techniques have emerged for studying RNA localization. Massively parallel reporter assays (MPRAs) enable the characterization of the localization of thousands of sequences at once and have been successfully used to identify localization elements active in mammalian neuronal cells from endogenous mRNA sequence [27–29]. With these approaches, previous work identified 260 nt sequences derived from the 3′ UTRs of several highly neurite-enriched mammalian RNAs that were necessary and sufficient for robust, kinesin-dependent RNA trafficking to neurites and more generally to the plus end of microtubules in a variety of cell types [27,30,31]. However, it remained unclear what sequences within or properties of these elements were endowing them with RNA localization activity. In comparing the LEs derived from different genes, there were no obvious characteristics or motifs shared among them. Using MPRAs to systematically test features and subsequences of these 260 nt LEs, we sought to identify the characteristics critical for their activity with high resolution.

Here, using MPRAs with over 50,000 unique, systematically designed reporter sequences based on previously identified LEs, we discover multipartite, highly active mammalian RNA localization elements that are approximately 200 nt in length. We further find that these LEs contain small regions requiring a strict sequence identity and larger regions that are tolerant to changes in nucleotide order but not nucleotide identity. These findings establish robust mammalian LEs as longer and more complex than sequence elements that regulate other facets of RNA metabolism [21].

## RESULTS

### Design of high-resolution MPRA to characterize RNA localization elements

Previous work from our laboratory identified a set of 260 nt-long RNA sequences that were necessary and sufficient for efficient kinesin-dependent RNA transport to the plus end of microtubules across a range of cell types [27,30,31], including to neurites in neuronal cells. These 260 nt sequences were identified within the 3′ UTRs of endogenous mRNAs which were also highly enriched in neurites [27]. However, it remained unclear what characteristics or parts of these 260 nt sequences were critical for RNA transport activity.

We therefore set out to determine exactly which parts of the 260 nt localization elements (LEs) were sufficient to drive RNA transport to neurites. To do this, we designed a massively parallel reporter assay (MPRA) in which the localization activities of all possible 5, 10, 20, 35, 50, 75, 100, 125, 150, 175, 200, 225, and 250 nt long contiguous subsets of the parent LEs were quantified (**Figure 1A**). We refer to this design as the “sufficiency MPRA”.

**Figure 1.**
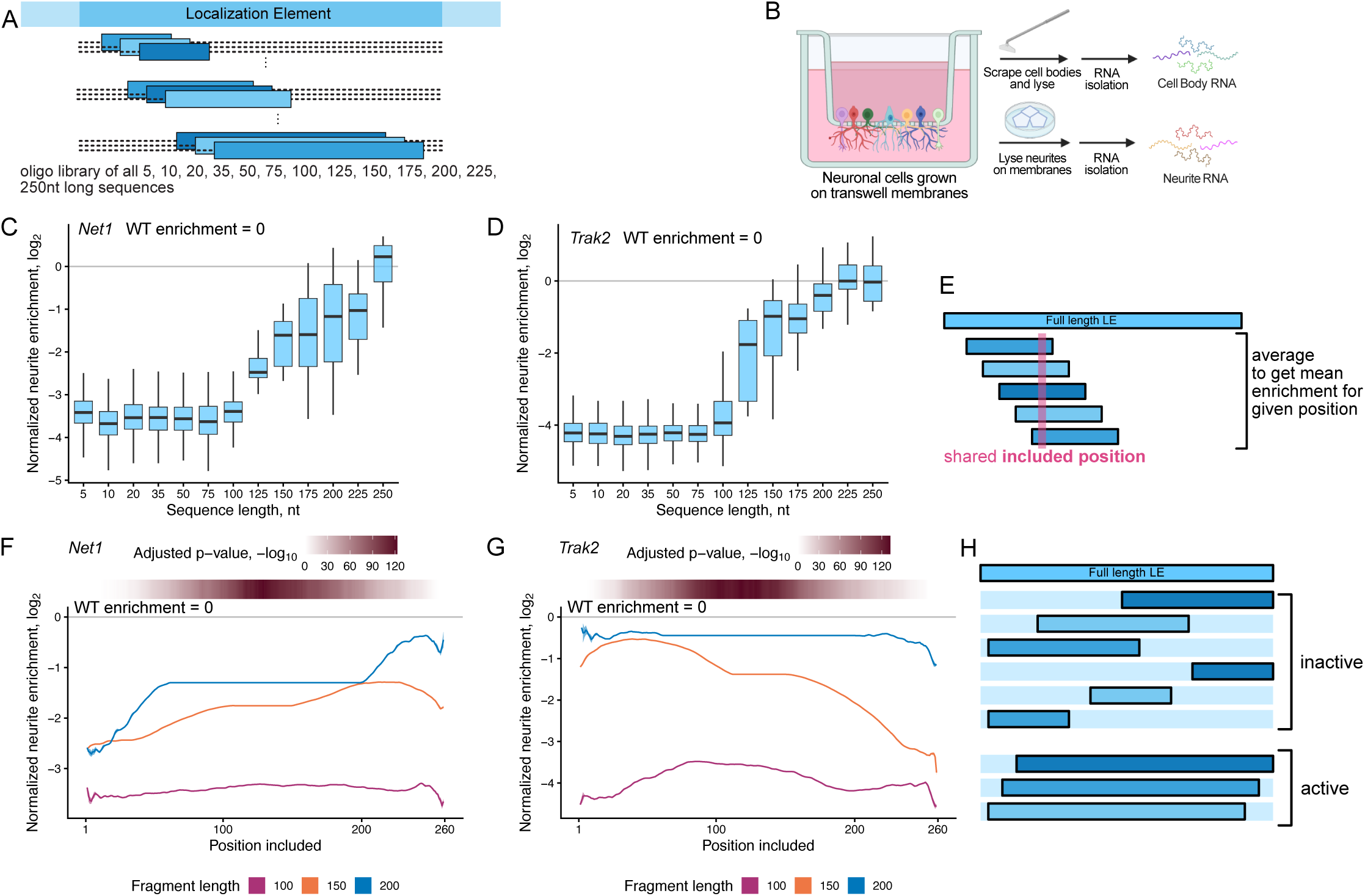
Mammalian neuronal localization elements have a minimum size greater than 200 nt. (A) Design of MPRA for uncovering the minimal size LE needed for RNA localization to neurites. (B) Schematic of subcellular fractionation and RNA isolation. (C) RNA localization to neurites based on the sequence subset length of the *Net1* LE tested relative to 250 nt fragments of *Net1* LE enrichment. (D) RNA localization to neurites based on the sequence subset length of the *Trak2* LE tested relative to 250 nt fragments *Trak2* LE enrichment. (E) Schematic of calculation of neurite RNA enrichment for each sequence subset containing a given position. (F) Neurite enrichment for reporters containing 100, 150, and 200 nt long *Net1* LE sequence subsets derived from the parent 260mer LE. Rolling mean, k=3. Ribbon represents standard error. Purple represents 100nt sequence subsets, orange represents 150 nt sequence subsets, and blue represents 200nt subsets. The flat section in the center of the 150 and 200 nt lines represents the mean enrichment of all sequence subsets of that size, as all possible subsets of that size contain those central positions. P-values were calculated for each position by using a Wilcoxon rank-sum test comparing all sequences derived from the *Net1* LE that contain a position to all those that do not. Power of this test is much greater at positions in the center of the LE as there are more oligos that contain those positions. P-values were adjusted using a Bonferroni correction. (G) As in F, but for *Trak2* LE. (H) Schematic of conclusions made from sufficiency MPRA.

In this MPRA, oligos containing these LE subsequences were synthesized and incorporated into the 3′ UTRs of two separate reporter RNAs that encoded GFP and Firefly luciferase. The resulting libraries of reporters were then site-specifically integrated into the genome of CAD mouse neuronal cells via Cre-mediated recombination [32]. We then mechanically fractionated these cells into soma and neurite fractions by growing them on porous transwell membranes [33] and isolated RNA from each fraction. We determined the relative abundance of every reporter-embedded oligo in these RNA fractions using targeted RT-PCR followed by high-throughput sequencing. Unique molecular identifiers incorporated during reverse transcription allowed us to accurately quantify reporter transcript abundances in RNA samples without the introduction of noise from PCR amplification. By comparing the relative abundances of reporter-embedded oligos in soma and neurite RNA fractions, we were able to quantify their relative neurite enrichments, *i.e.* their localization activities (**Figure 1B, Table S1, Supplementary File 1**).

For every oligo, this neurite enrichment was normalized to the neurite enrichment of the full, 260 nt wildtype (WT) LE. All oligos that were less neurite enriched than the parent wildtype LE therefore had a negative normalized neurite enrichment and those that were more neurite-enriched than their parent LE had positive normalized neurite enrichment values.

### The minimally active size of highly active mammalian LEs is approximately 200 nt

We tested LEs derived from seven different genes but chose to focus on those derived from the 3′ UTRs of *Net1* and *Trak2* as their LEs consistently drove approximately 20-fold RNA enrichment in neurites. In the cases of both of these LEs, oligos containing sequence subsets of 125 nt or smaller were 4- to 16-fold less neurite enriched than wildtype (**Figure 1C-D**). Sequence subsets larger than 125 nt drove approximately half of the wildtype localization activity, but only subsets above 225 nt for *Net1* (**Figure 1C**) and 200 nt in length for *Trak2* (**Figure 1D**) approached levels of neurite enrichment seen with the full-length, wildtype LE. These trends are maintained within the other fragment sizes (**Figure S1A-B**).

The relatively large minimum size for highly active sequence subsets of LEs is consistent with the sizes of the first localization elements defined in highly localized transcripts in *D. melanogaster* embryos, *Xenopus* oocytes, chicken embryonic fibroblasts, rat hippocampal primary culture cells, and *S. cerevisiae*. In these cases, the identified LEs were approximately 100-200 nt [1–4,9].

We then aimed at identifying regions within the 260 nt LEs that conferred RNA localization activity to reporter RNAs. To do so, for each nucleotide position within the 260mer, we plotted the mean neurite enrichment of all reporter RNAs whose embedded oligo contained that position (**Figure 1E**).

With this visualization, we see that 100 nt sequence subsets drawn from the 260 nt LEs of *Net1* and *Trak2* are again unable to drive localization activity, regardless of from where within the LE they are derived (**Figure 1F, G**). However, larger 150 nt subsets from both LEs can drive some neurite enrichment, depending on the location within the LE from which they were derived.

For the *Net1* LE, fragments that include the 3′ end of the LE have more neurite enrichment activity than those drawn from the 5′ end (**Figure 1F**). Conversely, in the *Trak2* LE, inclusion of the 5′ end of the LE led to greater neurite enrichment (**Figure 1G**). 200 nt fragments tended to show higher levels of localization activity than the 150 nt fragments with the same positional biases (**Figure 1F-G**).

From these data, we conclude that the minimum active size of these LEs is approximately 200 nt. Furthermore, specific regions within these LEs are more important for driving RNAs to neurites than others (**Figure 1H**).

### Small regions within LEs are critical for RNA localization activity

Having defined the minimally active size of LEs to be approximately 200 nt via the sufficiency MPRA, we next wondered if specific smaller regions of the LEs were necessary for activity. To interrogate this, we designed an MPRA where we made deletions within sliding windows along the LEs. We designed oligos that deleted all possible 5, 10, 20, 50, 100, 150, and 250 nt contiguous windows across the LEs (**Figure 2A**) and again incorporated them into the 3′ UTRs of reporter transcripts. In this way, we test the necessity of each window for RNA localization activity. We refer to this design as the “necessity MPRA.”

**Figure 2.**
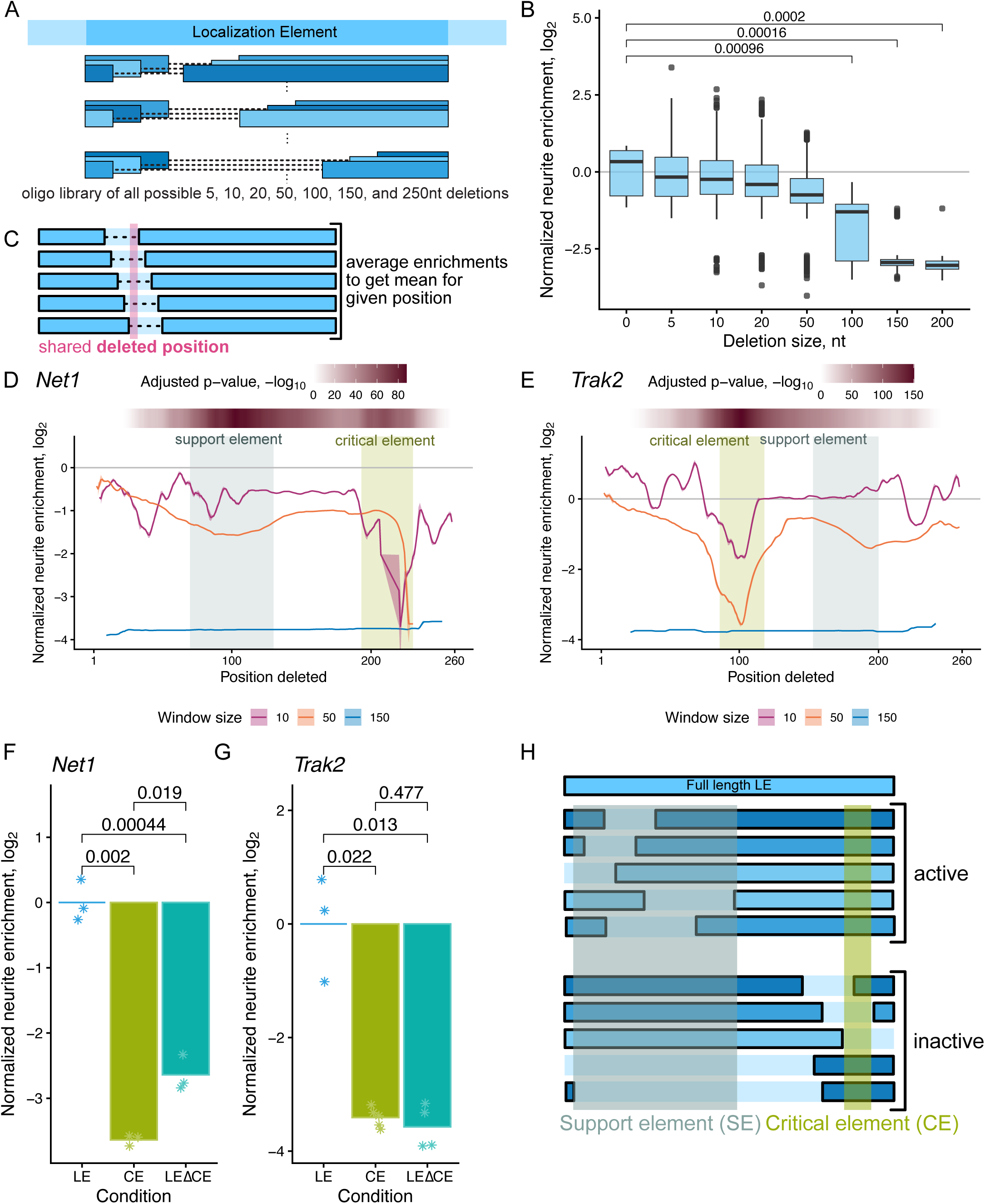
LEs contain two subelements, including a necessary critical element. (A) Design of MPRA for finding necessary elements of LE for RNA localization to neurites. (B) RNA localization to neurites of all LEs depending on deletion window size. P values were calculated using a Wilcoxon rank-sum test. Non-significant values not shown for differences between 0 and other deletion sizes. (C) Schematic of calculation of neurite RNA enrichment for each oligo missing a given position. (D) Neurite enrichment for reporters containing *Net1* LE derived sequences containing deletion window sizes of 10, 50, and 150 nt. Rolling mean, k=3. Ribbon represents standard error. Purple represents 10nt window deleted, orange represents 50nt window deleted, and blue represents 150nt window deleted. CE is indicated by the light green background. SE is indicated by the blue-grey background. P-values were calculated for each position by using a Wilcoxon rank-sum test comparing all sequences derived from the *Net1* LE that have a position deleted to all those that do not. Power of this test is diminished at the ends of the LE as fewer oligos have those positions deleted. P-values were adjusted using a Bonferroni correction. (E) As in D, but for *Trak2* LE. (F) The indicated sequences derived from the *Net1* LE were fused to the 3′ UTR of firefly luciferase and the neurite localization of the resulting transcript was assayed using RT-qPCR. P-values were calculated using Student’s T-test. (G) As in F, but for *Trak2* LE. (H) Schematic of conclusions made from necessity MPRA.

As expected, the wildtype LEs, with deletion size zero, had the greatest neurite enrichment. As the deletion sizes within the oligos tested increased, the neurite localization decreased. Generally, the deletion of small 5 or 10 nt windows was tolerated, while the deletion of larger windows led to larger decreases in RNA neurite enrichment (**Figure 2B, Table S2, Supplementary File 1**). However, specific smaller deletions resulted in ablation of localization activity to levels similar to those seen for larger deletions (**Figure 2B**). This suggests that LEs may contain small sequence windows that are critical contributors to their localization activity (**Figure 2B**).

To determine where these small sequence windows could be, we plotted the average neurite enrichment of all reporter RNAs whose embedded oligo had a given position deleted for each position within the 260 nt LE (**Figure 2C**). This is similar to how we plotted the sufficiency MPRA, but to examine necessity, we are averaging together reporters whose embedded oligo does not contain a given position in the wildtype sequence.

For *Net1*, most 10 nt deletions resulted in only a slight decrease of localization activity compared to wildtype. However, if the 10 nt deletion window resulted in the removal of any of positions 193-230 of the 260 nt LE, it resulted in a large decrease in RNA localization activity (**Figure 2D)**. Similar patterns were observed with the *Trak2* LE, where 10 nt deletions were generally well tolerated across the entire LE, except when any of positions 86-118 were removed (**Figure 2E**). These effects became more pronounced at larger deletion sizes. In both the *Net1* and *Trak2* LEs, deletions of 150 nt resulted in oligos without localization activity.

### Critical elements are necessary but not sufficient to drive RNA localization

We refer to the small regions whose deletion resulted in large decreases in RNA localization activity in the necessity MPRA as “critical elements” (CEs) (**Figure 2D-E**). To more thoroughly test whether the CEs were necessary and sufficient for localization activity, we designed reporters containing either the full-length LE, a region overlapping the CE, or the full-length LE minus that section (*Net1* LEΔCE, *Trak2* LEΔCE).

These reporters were expressed in CAD cells, and their neurite enrichment was assessed via subcellular fractionation and RT-qPCR. In the case of both *Net1* and *Trak2*, their CEs alone failed to drive localization of the reporters to the neurites. When only the CE was removed from the wildtype LE, the remaining sequence was also unable to drive localization (**Figure 2F-G**). These data indicate that the CEs are necessary but not sufficient for RNA transport.

### Support elements, which are separate from critical elements, also contribute to localization activity

Although small deletions that overlapped the CEs resulted in drastic reductions in RNA localization activity, larger, 50 nt deletions in other regions of the LE that did not overlap with the CE also decreased activity. In the *Net1* LE, 50 nt deletions within positions 70-130 diminished neurite localization (**Figure 2D**), and in the *Trak2* LE, 50 nt deletions between positions 153-200 also led to diminished activity (**Figure 2E**). Notably, we know from the sufficiency MPRA (**Figure 1**) that these regions alone cannot drive neurite RNA enrichment. We therefore refer to these elements as “support elements” (SEs). Interestingly, SEs generally tolerate small deletions up to 20 nt, but their contributions to localization activity decrease with deletions 50 nt or larger (**Figure S2A-B**).

Together, this data allowed us to revise our model. Sequences that include both the entire CE and a large portion of the SE are most active. Small deletions to the SE are tolerated, but small deletions of the CE are not (**Figure 2H**).

### Critical elements are necessary for axonal RNA localization in primary neurons

To further explore the biological relevance of the CEs, we then probed their activity in primary rat hippocampal neurons. We reasoned that since we had previously observed that *Net1* and other RNAs are strongly trafficked to the plus end of microtubules, CEs within them might be crucial for trafficking to axons [27,30,31].

To test this, we used reporter RNAs containing a lacZ open reading frame followed by 128 MS2 hairpins in the 3′ UTR that allow visualization of single RNA molecules in cells via co-expressed fluorescent MS2 coat protein [34]. We then incorporated LEs from *Net1* and *Trak2* that were either wildtype or contained CE deletions into the 3′ UTRs of these reporters. We expressed these constructs in rat hippocampal neurons and measured the subcellular distribution of reporter RNAs using fluorescence microscopy.

For each construct, we quantified the number of fluorescent puncta present in Tau-positive axons (**Figure 3A**). For both *Net1* and *Trak2*, we found that loss of the CE resulted in a significant decrease in reporter RNA abundance in axons, suggesting its necessity for axonal transport (**Figure 3B**). Importantly, these results were not due to the wildtype LEs simply leading to higher overall expression. In fact, wildtype reporter RNAs were less abundant in dendrites than their CE-lacking counterparts (**Figure S3A**), potentially due to wildtype RNAs being specifically targeted to axons. Given that CAD neurites are primarily axonal in character [30,35], these findings indicate that our MPRA results in neuronal cell lines likely faithfully recapitulate trafficking patterns in primary cells.

**Figure 3.**
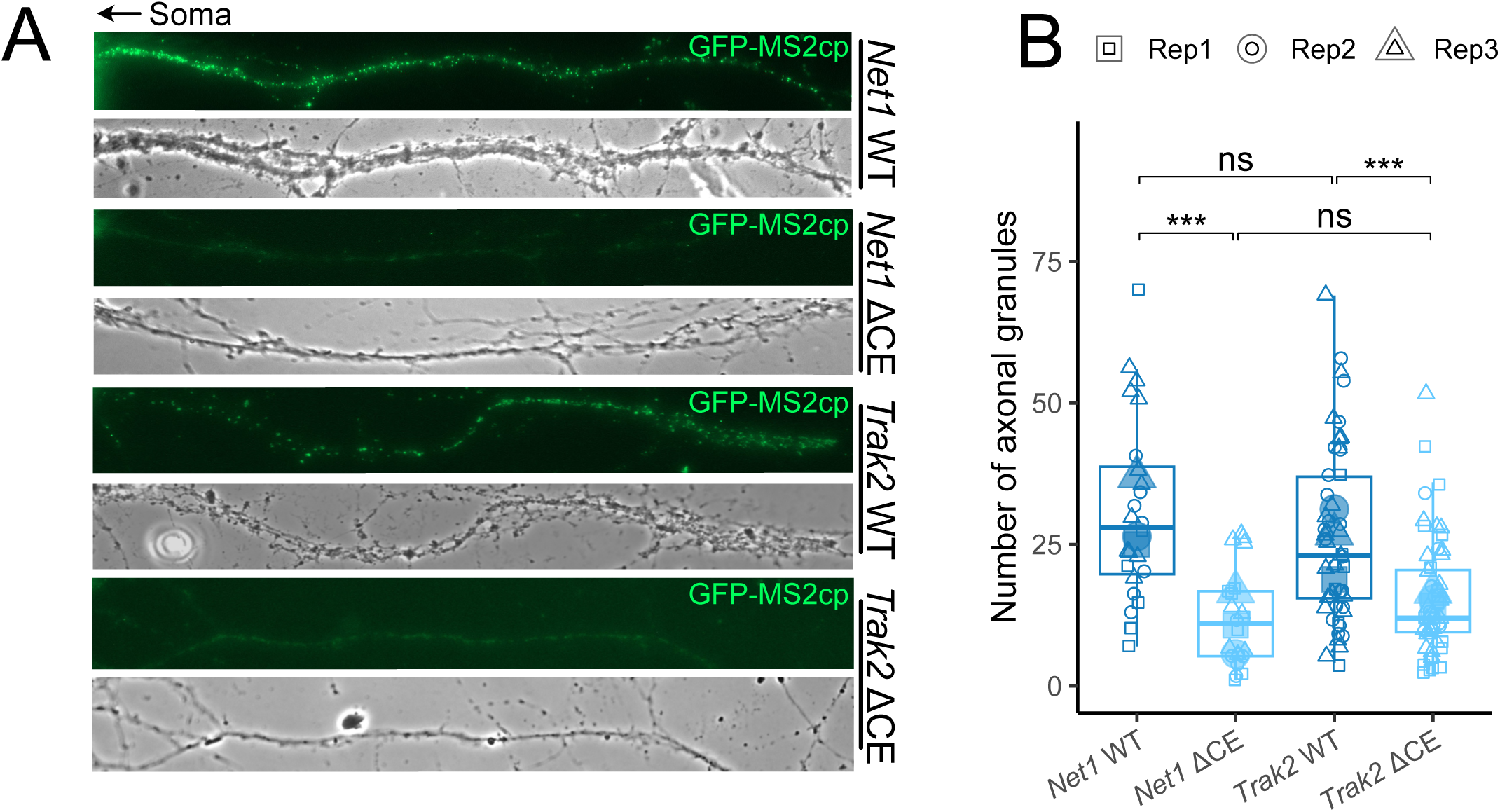
Transport of LE RNAs to hippocampal neuron axons depends on the CE. (A) Representative images of hippocampal neuron axons expressing MS2 tagged LE constructs (green). (B) Quantification of the number of LE construct-containing granules in axons.

### Mutating sequences within critical elements greatly reduces localization activity

Thus far, all of our tested reporters were composed of elements where sections of the parental, active LE had been removed, thus altering the spacing between sequences within the elements. To preserve the spacing between elements but still interrogate their contribution to localization activity, we designed another MRPA using a similar approach to the previous, deletion-based, necessity MPRA. Instead of deleting the sequences within each possible 5, 10, 50, 100, and 200 nt window, we randomly mutated the sequence within that window. This mutation was done three times per window to create three unique sequences per window tested (**Figure 4A**).

**Figure 4.**
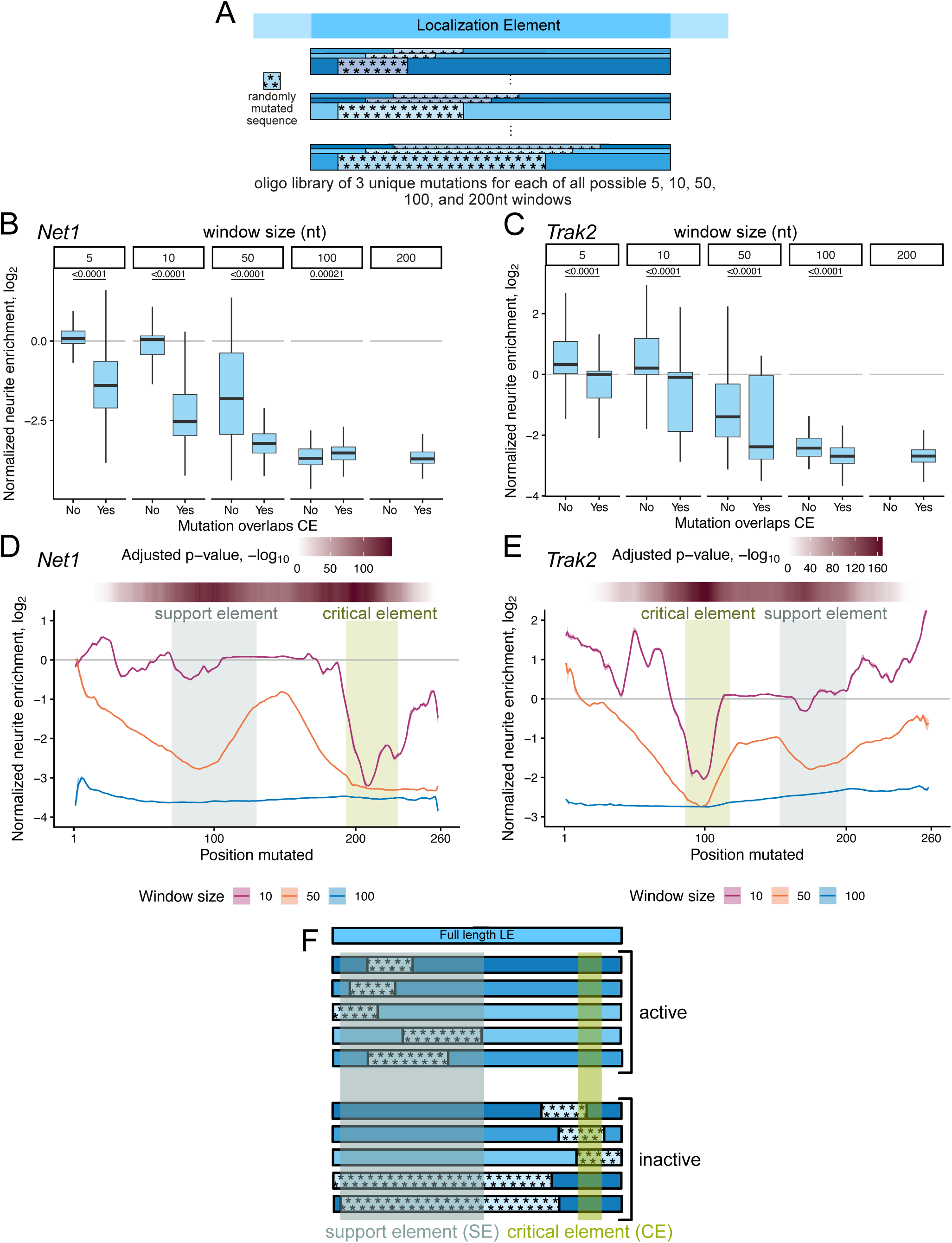
Support elements tolerate small mutations, but critical elements are intolerant to change. (A) Design of MPRA for understanding the influence of sequence identity on RNA localization. (B) Comparing RNA localization to neurites of reporters that contain mutation windows of sizes 5, 10, 50, 100, and 200 nt and do or do not mutate any part of the *Net1* CE. Due to the positioning of the *Net1* CE, there are no possible 200 nt windows that do not overlap with the CE. P-values were calculated using a Wilcoxon rank-sum test. (C) As in B, but for *Trak2* LE. Due to the positioning of the *Trak2* CE, there are no possible 200 nt windows that do not overlap with the CE. (D) Neurite enrichment for reporters containing *Net1* LE derived sequences containing mutation window sizes of 10, 50, and 100 nt. Rolling mean, k=3. Ribbon represents standard error. Purple represents 10nt window mutated, orange represents 50nt window mutated, and blue represents 100 nt window mutated. CE is indicated by the light green background. SE is indicated by the blue-grey background. P-values were calculated for each position by using a Wilcoxon rank-sum test comparing all sequences derived from the *Net1* LE that have a position mutated to all those that do not. Power of this test is diminished at the ends of the LE as fewer oligos have those positions mutated. P-values were adjusted using a Bonferroni correction. (E) As in D, but for *Trak2* LE. (F) Schematic of conclusions made from mutation MPRA.

Given that the necessity MPRA demonstrated the sensitivity of the CEs to deletion, we first examined the effect of mutations that changed at least one position within the CEs. Consistent with previous results, making even small mutations to the CE significantly reduced localization activity (**Figure 4B-C, Table S3, Supplementary File 1**) while small mutations outside the CE were better tolerated (**Figure S4A-C**). Larger mutation windows were not tolerated regardless of if they overlapped the CE or not (**Figure 4B-C**).

### Support elements tolerate small mutations

When further examining how mutation of the LE affects its activity, we found that SEs tolerated smaller, 10 nt wide mutation windows, but not larger 50 or 100 nt mutation windows (**Figure 4D-E, S4B-C**). These results from the mutation MPRA mirror what was seen with the deletion of the same sequences in the necessity MPRA. From this, we conclude that both the presence of sequence at the CEs and SEs and the identity of those sequences are important for RNA localization activity (**Figure 4F**).

### The identity, but not the order, of nucleotides within support elements is critical for activity

Sequence identity, as tested in the mutation MPRA (**Figure 4**), is an important factor for determining the activity of a LE. However, the overall content of each nucleotide (e.g. proportion of adenosines, proportion of uridines, etc.) within an element could be more important than the exact order of those nucleotides. For example, the sequence contents of RNA elements have been found to be correlated with RNA localization, with nuclear-retained RNAs being more C/U rich and neurite localized RNAs being more A/G rich [27,36,37].

To determine the effect that overall sequence content, rather than specific nucleotide order, has on LE activity, we designed another MPRA, which we termed the “shuffle MPRA”. Similar to the mutation MPRA, we made changes within all possible 5, 10, 50, 100, 200, and 225 nt windows. This time, however, instead of altering the identities of nucleotides within a given window, we simply shuffled their order within the window (**Figure 5A**). To ensure that we tested divergent sequences, we required that the identities of the bases at a given position be changed in at least half of the positions within the window when possible. As seen in the necessity and mutation MPRAs, the larger the size of the window perturbed, the less neurite RNA enrichment activity the resulting sequences displayed (**Figure 5B, S5A, Table S4, Supplementary File 1**). Surprisingly, within a given window size, shuffles that resulted in more position-specific nucleotide identity changes did not reliably result in larger decreases in localization activity (**Figure 5B, S5A**).

**Figure 5.**
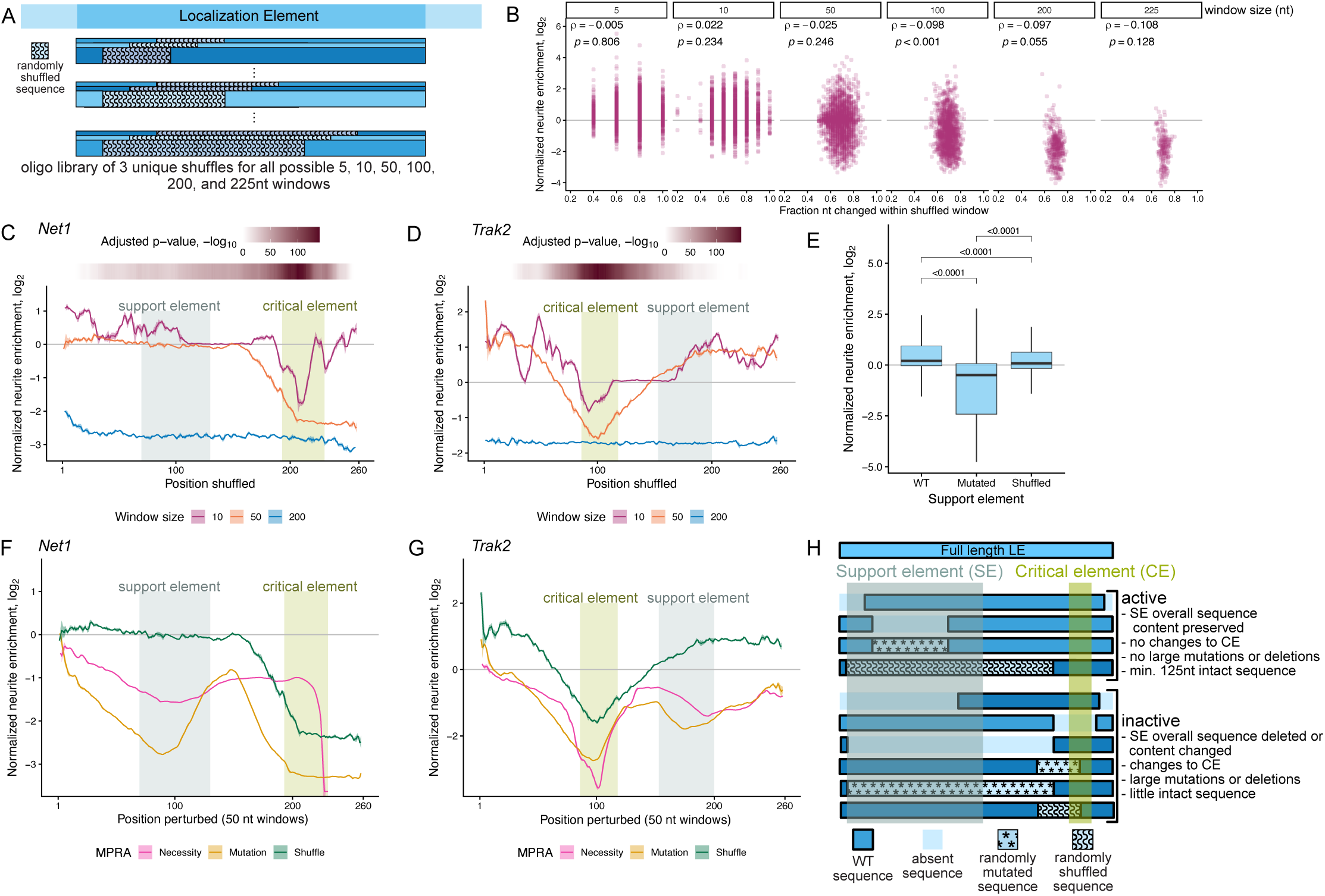
Support element localization activity depends on sequence composition. (A) Design of MPRA for understanding the influence of overall sequence content on RNA localization. (B) Neurite enrichment of reporters from all LEs depending on the size of the window shuffled and the percentage of nucleotide identities within the changed. Vertical lines in the plots for smaller window sizes are due to the small number of positions to change (i.e. for a 5 nt window changing the identity of 1 position means 20% of the sequence changed, changing the identity of 2 means 40% of the sequence is changed ect.). Correlation coefficient calculated using the Spearman method. (C) Neurite enrichment for reporters containing *Net1* LE derived sequences containing shuffle window sizes of 10, 50, and 200 nt. Rolling mean, k=3. Ribbon represents standard error. Purple represents 10 nt window shuffled, orange represents 50 nt window shuffled, and blue represents 200 nt window shuffled. CE is indicated by the light green background. SE is indicated by the blue-grey background. P-values were calculated for each position by using a Wilcoxon rank-sum test comparing all sequences derived from the *Net1* LE that have a position within the shuffled window to all those that do not. Power of this test is diminished at the ends of the LE as fewer oligos have those positions within a shuffled window. P-values were adjusted using a Bonferroni correction. (D) As in C, but for *Trak2* LE. (E) Neurite enrichment of reporters derived from the *Net1* and *Trak2* LEs with wildtype, mutated, and shuffled SEs. P-values were calculated using a Wilcoxon rank-sum test. (F) Neurite enrichment for reporters containing *Net1* LE derived sequences that have been subjected to a 50nt wide window of perturbation. Purple represents reporters from the necessity MPRA where 50 nt windows have been deleted. Orange represents reporters from the mutation MPRA where 50 nt windows have been randomly mutated. Blue represents reporters from the shuffle MRPA where 5 nt windows have been randomly shuffled. CE is indicated by the light green background. SE is indicated by the blue-grey background. (G) As in F, but for *Trak2* LE. (H) Summary schematic of conclusions collected from all MPRAs.

When examining how the location within the LE of the shuffled region affected RNA localization activity, we again found that for both *Net1* and *Trak2*, even small shuffles that overlapped the CEs were associated with a large decrease in activity (**Figure 5C-D**). This is consistent with results from the necessity and mutation MPRAs in that CEs do not tolerate any changes. Although shuffles of 200 nt windows were not tolerated in any regions, 50 nt shuffles that overlapped the SEs were generally tolerated (**Figure 5C-D, S5B-C**).

To interrogate this more thoroughly, we directly compared the results of mutating and shuffling nucleotides within the SE across all tested sequences from the mutation and shuffle MPRAs. We found that mutation of sequences within the SE had a much larger negative effect on RNA localization activity than shuffling the order of the nucleotides within the SE (**Figure 5E**). This suggests that the overall sequence content (i.e. the fraction of each nucleotide within a sequence), rather than the exact order of nucleotides, of the SE defines the majority of its contribution to RNA localization activity.

These results are also seen when comparing the results of the necessity, mutation, and shuffle MPRAs (**Figure 5F-G**). Across these MPRAs, 50 nt windows of perturbation that overlap the CE result in large decreases in RNA localization activity, regardless of the type of perturbation. However, with the SE, only deletion or mutation of 50 nt windows resulted in decreased activity. Shuffling the order of nucleotides within the exact same window had no effect on RNA localization activity.

Based on these findings, we updated our model of LE function (**Figure 5H**). Any type of perturbation to the CE results in a loss of function. Mutation or deletion of the SE similarly results in a loss of function, but random shuffling of the nucleotides within the SE does not. This supports the idea that the contribution of the SE to RNA localization activity depends on its overall sequence content rather than on the exact order of its nucleotides.

### Investigation of the role of RNA secondary structure in LE activity

Many LEs identified in other transcripts, particularly in *Drosophila*, require specific secondary structures for their function [20,23,38–42]. As such, we sought to determine the role of RNA secondary structure in the activity of the *Trak2* and *Net1* LEs. To determine the possible secondary structure of these sequences, we used 1M7 (1-Methyl-7-nitroisatoic anhydride) to chemically probe the LEs in solution using SHAPE-MaP [43]. In the SHAPE-informed minimum free energy model, the *Net1* LE formed a 5-stem junction and a stem loop (**Figure 6A**), with many structural features supported by high pairing probabilities from the partition function (**Figure S6A**). The *Net1* CE contained the most 3′ double stranded region of the junction and the most 3′ stem loop. The SE contained the 3 most 5′ stems of the 5-stem junction (**Figure 6A**). The *Trak2* LE formed two stem loops with a single branch separated by a single stranded region (**Figure 6B**), although SHAPE-informed modeling suggests that this RNA is more intrinsically dynamic, with fewer of the minimum free energy structures highly populating the ensemble (**Figure S6B**). The *Trak2* CE spanned the 3′ end of the first stem loop and the single stranded region. The SE was found at the most distal loop of the 3′ branched stem loop (**Figure 6B**).

**Figure 6.**
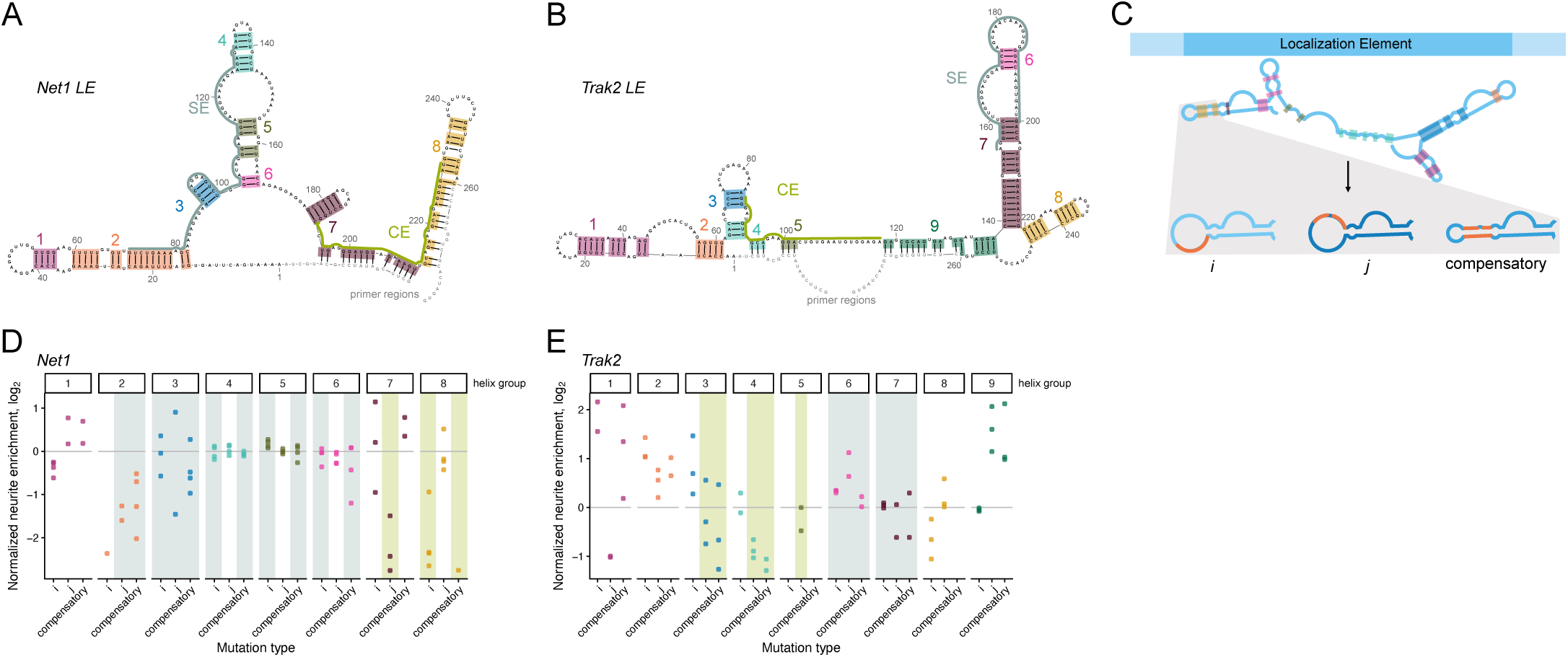
SHAPE-informed secondary structures of LEs do not confer localization activity. (A) SHAPE-determined structure of the *Net1* LE, including primer sequence (grey). Groups of double stranded sequences are highlighted and numbered in corresponding colors (group 1 - purple, group 2 - orange, group 3 - blue, group 4 - turquoise, group 5 - olive, group 6 - pink, group 7 - dark purple, group 8 - gold). CE is indicated with a light green line along the sequence in the structure. The SE is similarly indicated with a blue-grey line. (B) As in A, but for the *Trak2* LE. Groups of double stranded sequences are highlighted and numbered in corresponding colors (group 1 - purple, group 2 - orange, group 3 - blue, group 4 - turquoise, group 5 - olive, group 6 - pink, group 7 - dark purple, group 8 - gold, group 9 - green). (C) Design of MPRA for testing the functionality of SHAPE-determined structures of LEs. (D) Neurite enrichment of *Net1* reporters with a single group of double stranded regions mutated. i mutations are made to the 5′ strand of the group, j mutations to the 3′ strand, and compensatory mutations made to both stands to create different sequences that restores stem formation. The background shading indicates which subelement is affected by mutations made to that helix group. Blue-grey shading indicates the SE is mutated and green shading indicates that the CE is mutated. (E) As in D, but for *Trak2* LE. Group 5 only has j mutations as i or compensatory mutations would disrupt the sequence necessary for sequencing library amplification.

To test if the secondary structures that we determined contributed to the localization of the LEs, we grouped the double stranded regions based on the number of unpaired bases predicted to interrupt each paired region (**Figure 6A-B**). We then designed another MPRA, termed the SHAPE MPRA. In this MPRA, we made random mutations to the bases predicted to be paired. We independently mutated the 5′ (i) and 3′ (j) sides of each helix group. Then, we created a separate group of oligos that contained compensatory mutations that changed both sides of a helix group but maintained the base-pairing potential between the i and j bases. This was done for all possible combinations of helix groups (**Figure 6C, Table S5, Supplementary File 1**).

For the *Net1* LE, mutations of any type to helix groups 4, 5, 6, and 7 did not affect localization activity (**Figure 6D**). These mutations at most change 10 nt within the SE, which preserves its function as seen in the mutation MPRA (**Figures 4D, 5C**). Stem j mutations to group 7 and i and compensatory bases of group 8 greatly disrupt localization activity, which is in line with previous MPRAs, as those changes disrupt the CE. Mutations to group 1 do not substantially change neurite enrichment (**Figure 6D**). This is in line with previous results where small mutations in the most 5′ end of the *Net1* LE minimally disrupt localization (**Figures 4D, 5C**). Mutations to group 2 do markedly decrease localization as those changes are larger, again in line with previous observations in the mutation MPRA (**Figures 4D, 5C, 6D**). However, in no helix group did we observe a loss of activity in i and j mutations that was restored by the compensatory mutations.

In general, we observed similar trends with the *Trak2* LE. Mutations to helix groups 6 and 7, both small regions of the SE, did not result in significant decreases in activity. Conversely, the j mutations in helix groups 3, 4, and 5 as well as the compensatory mutations in groups 3 and 4 disrupt the CE and are associated with larger decreases in activity (**Figures 6B, 6E**). This is again consistent with the results from the mutation and necessity MPRAs. As with *Net1*, we did not observe any cases where localization activity losses due to mutations in both i and j stems were restored by compensatory mutations (**Figure 6E**). These results suggest that although secondary structure may be involved in the activity of these localization elements, the structures we determined do not contribute to activity.

## DISCUSSION

While RNA localization is known to be critical for neuronal function, it remains unclear what makes a given RNA a target for neurite localization [7,8,10–14]. Prior work in defining the *cis*-elements needed for RNA localization has necessitated serially testing constructs to refine what sequences constitute the minimal LE [3,9,23,44–47]. With the exceptions of classically studied RNAs like those encoding beta-actin and myelin basic protein, we do not adequately understand what sequences are needed for localization to neuronal cell projections [3,48–51]. Previous work from our lab addressed this problem using MPRA approaches to identify 260 nt sequences within the 3′ UTRs of several transcripts that were necessary and sufficient for transport to neurites. However, it remained unclear what features within those 260 nt sequences enabled their ability to drive RNA transport or what subsequences within them that were critical for activity.

Here, we interrogate these elements further by testing the effects of systematic mutations, truncations and deletions on their ability to regulate RNA localization in mammalian cells. To do this, we used a series of MPRAs, allowing us to test the localization activity of approximately 50,000 unique sequences derived from the original 260 nt LEs [27]. We determined that the minimum size for robust neurite localization is approximately 200 nt. This is consistent with previous *Drosophila* literature on localization-regulatory *cis*-elements [38,44,46,52].

It is also consistent with previous MPRA-based studies in mammalian neurons [28,29]. These studies used smaller UTR fragments, usually 150 nt, in their assays, thus rendering them unable to identify larger elements. Generally, these studies found elements that drove RNAs to approximately 5-fold neurite enrichment. In contrast, the elements we identified using longer UTR fragments are capable of driving neurite enrichments of up to 20-30 fold.

Within these relatively large sequences, we found that LEs derived from *Net1* and *Trak2* required the activity of two subregions. The first, named the critical element or CE, was smaller and was highly intolerant of any changes to its sequence. The second, named the support element or SE, was larger and only required the majority of its sequence content to be intact to contribute to neurite localization. Interestingly, the precise order of the nucleotides within the SE was less important than their identities, suggesting that the overall sequence content of the SE is important for its function. We then determined the secondary structure of these LEs by SHAPE, which when functionally tested proved to not contribute to localization activity. While the SHAPE-informed secondary structures we examined proved to be non-functional, this does not exclude secondary structure from playing a role in the localization of these RNA sequences. Our non-functional secondary structure could be explained by these LEs folding differently in cells compared to *in vitro*, potentially due to interactions with a trans-factor. Alternatively, we could have been unable to detect the structure contributing to localization. For example, G-quadruplexes are known to contribute to RNA localization, but are not able to be detected by standard SHAPE-MaP [43,53].

Our study uncovered that mammalian LEs needed for neurite localization are highly complex and require multipartite motifs that depend on both primary sequences and overall nucleotide composition. The similarity in requirements yet considerable divergence in primary sequence between the LEs in *Trak2* and *Net1* raise the question of whether these two LEs are localized via the same mechanism. Further work will be required to determine if that is the case.

## ACKNOWLEDGEMENTS

We thank members of the Taliaferro lab for helpful discussions. This work was funded by NIH grants R35GM133385 (J.M.T.), R01NS122911 (J.M.T.), F31GM151819 (C.M.), T32GM136444 (C.M.), R01GM155542 (C.A.W.), and R35GM156773 (A.W.M.), by the DFG (M.A.K., Ki 502/9-1, 506658941) and the Friedrich Baur Foundation (K.E.B.).

## METHODS

### MPRA design

The code for designing all MPRA oligo pools is available at: https://github.com/TaliaferroLab/peak-oligo/blob/ master/united_MPRA_design.qmd

The sufficiency, necessity, mutation, and shuffle MPRAs are designed using a “sliding window” approach, where windows of various sizes are tiled across the parent 260mer LE at a step size of 1 nt. In the case of the sufficiency MPRA, the sequence within the window is preserved, then inactive sequence from the endogenous locus of parent LE is appended to the sequence window to create an oligo that is 260 nt long. This process is inverted for the necessity MPRA, where sequence within the window is removed. The remaining LE sequence is then padded to 260 nt with the same inactive sequence used for the sufficiency MPRA. The mutation and shuffle MPRAs have the sequence within the window randomly mutated or randomly reordered, respectively. To ensure divergent sequences were tested, we prioritized creating shuffles with a Hamming distance (i.e. the number of positions in the child sequence that are different than the parent sequence) greater than or equal to half the window size rounded up from the parent LE when possible.

The SHAPE MPRA is designed to mutate the 3′ and 5′ sides of a SHAPE-defined double stranded region to disrupt basepairing, then to mutate both to create a different sequence that restores stem formation (i.e. compensatory mutations). Double stranded regions were grouped into “stem” regions. Each “stem” is defined as a run of paired bases, interrupted by no more than 2 unpaired bases on the side with more unpaired bases during a break. This means that “stems” are not interrupted by bulges, but are interrupted by symmetric internal loops with 2 nt on each side, or asymmetric internal loops with a minimum of three nt on the shorter side. All possible combinations of “stems” were then mutated: the 3′ side (i), the 5′ side (j), and then compensatory mutations made (ij). In these oligos, no unpaired bases were changed. Each set of mutations was done in triplicate. The terminal loops of hairpin structures were manually annotated and all possible combinations were mutated five times. In these oligos, no paired bases were mutated. Similarly, all internal single stranded regions longer than 4nt were manually annotated and all possible combinations were mutated five times.

For all MPRAs, pools of oligonucleotides were synthesized by Twist Biosciences.

### Cloning of MPRA library

For each MPRA, as in [27], oligonucleotide pools were resuspended in 10 mM Tris–EDTA buffer, pH 8.0 to a concentration of 10 ng/µl. The 10ng of the pool was amplified by performing 8 PCR reactions 50 µl each, using KOD Hot Start Master Mix (Sigma-Aldrich, 71842) according to the manufacturer’s instructions for 15 cycles total. The oligonucleotide pool was amplified using primers specific to the 20-nt common sequence and an overhang containing sequence specific to the cloning site for each reporter. After amplification, the PCR reaction was digested with Exonuclease I, at 37°C for 2 h to digest the single-stranded template and primers. The DNA was then purified using 0.6× Nucleomag beads were used to get rid of longer DNA products. The supernatant from this purification was then removed and additional Nucleomag beads were added to bring the final overall concentration to 1.0×.

The reporter plasmid (pRD-RIPE, [32]) was linearized by digesting it with BstIX (NEB) at 37°C for 6 hours to clone the library into the 3′me UTR of the GFP reporter. Digested plasmid DNA was gel extracted using Zymoclean Gel DNA Recovery Kit (Zymo Research, #D4008). The digested plasmid and amplified DNA library were assembled using Gibson Assembly reaction (NEB). The insert:vector molar ratios used were: necessity-6:1, sufficiency-5:1, mutation-6:1, shuffle-9:1, SHAPE-5:1. The Gibson Assembly reaction was then incubated at 50°C for 30 min. The cloned reporter plasmid (∼200 ng DNA) was purified by ethanol precipitation to get rid of excess salts and was then transformed into *Escherichia coli* 10-beta cells (NEB, C3040) using a Biorad GenePulser electroporator. The transformed cells were grown in recovery medium at 37 °C for an hour and then plated on pre-warmed Luria broth (LB) agar-Carbenicillin 15-cm plates and incubated at 37 °C overnight. The next day, the colonies were harvested from the plates using LB medium and a spreader. The bacterial culture was then centrifuged at 4000 rpm for 20 min. The reporter plasmid libraries were purified using ZymoPURE Plasmid Maxiprep kit (Zymo Research, #D4203). Restriction digestion was performed to confirm that the plasmid library contains only a single insert of the right size.

The resulting plasmid libraries were the linearized by digestion with PmeI at 37°C for 6 hours to clone the library into the 3′ UTR of the Firefly reporter. The steps for cloning into the GFP 3′ UTR were repeated, using the following insert:vector molar ratios: FF: necessity-6:1, sufficiency-7:1, mutation-9:1, shuffle-5:1. SHAPE-7:1.

### Cell culture

CAD cells were grown in DMEM/F-12 (Gibco, #11320-033) supplemented with 10% Equafetal (Atlas Biologicals, #EF-0500-A) and 1% Penicillin–Streptomycin solution. The cells were grown in a humidified incubator at 37°C and 5% CO_2_.

### Genomic integration of MPRA library

CAD cells were plated in 6-well plates at 5 × 10e5 cells per well in respective growth medium 12–18 h before transfection. Cells were then co-transfected with the cloned reporter plasmid mixed with 1% of plasmid expressing Cre recombinase. To transfect one well of a 6-well plate, 1.5 µg of reporter plasmid and 15 ng of Cre-plasmid was mixed with 3 µl Lipofectamine LTX reagent (Invitrogen, #15338100), 1.5 µl PLUS reagent and 100 µl Opti-MEM following the manufacturer’s protocol. 8 6-well plates were transfected for the ∼12,000 oligo MPRAs (SHAPE, necessity, sufficiency) and 12 6-well plates were transfected for the ∼18,000 oligo MPRAs (mutation, shuffle).

Cells were incubated with the transfection mixtures for 24 h, followed by the media change. The cells were incubated for an additional 24 h allowing for recovery and expression of the antibiotic resistance before addition of puromycin (5 µg/mL). The cells were selected in the puromycin until the cells in the control wells died. The cells with stably integrated reporter plasmids were expanded in the growth medium with 2.5 µg/mL puromycin.

### Subcellular fractionation

Expression of reporters for a given MPRA was induced by adding 1 µg/mL doxycycline to complete growth medium. In parallel, the same was done to cells expressing the MPRA from [27]. 48 hours later, both sets of MPRA expressing cells were mixed 2 parts of the experimental MPRA to 1 part of the MPRA from [27]. Cells were plated on transwell membranes and serum starved to differentiate, while being maintained in 1 µg/mL doxycycline. 48h later, cells were fractionated as in [27].

To ensure the quality of the fractionation, the 25ng of RNA from both the soma and neurite fractions of a single replicate were subjected to RT-qPCR (see below) to assay the levels of endogenous *Net1* and *Tsc1* mRNA, and to thus calculate the neurite enrichment of endogenous *Net1*. As endogenous *Net1* mRNA is known to be 20-30 fold neurite enriched, soma-neurite sample pairs were subjected to library preparation and sequencing only if they exhibited an endogenous *Net1* neurite enrichment of 16 fold by RT-qPCR. [27]

### MPRA sequencing library preparation

As in [27], 500 ng total RNA from each soma and neurite fractions was taken to synthesized cDNA in a 20 µl reaction using SuperScript IV Reverse Transcriptase (ThermoFisher) according to the manufacturer’s protocol, with primers specific to Firefly and GFP CDS containing an 8-nt unique molecular identifier (UMI) and a partial Illumina read 1 primer sequence. The incubation time at extension step (55°C) was increased to an hour. Post reverse transcription, 1µL each of RNAse H and RNaseA/T1 mix was added directly into the RT-reaction and incubated at 37°C for 30 min to digest the remaining RNA and RNA:DNA hybrids. The cDNA was purified using Zymo DNA Clean & Concentrator kit (Zymo, #D4013) using 7:1 excess of DNA binding buffer recommended for binding ssDNA.

For library preparation, each purified reporter cDNA reaction was split into five PCR reactions (4 µL cDNA/ PCR) and amplified using a reporter specific forward primer with Illumina sequencing adaptors and a reverse primer binding the partial Illumina read 1 sequence with the remaining sequencing adaptors using Kapa HiFi HotStart DNA Polymerase (Kapa Biosystems, #KK2601) using 18× cycles for GFP and 23× for Firefly reporter. The five PCR reactions per sample were pooled together and purified using double Nucelomag beads protocol.

In the first purification round, 0.6× Nucleomag beads were used to get rid of longer DNA products. The supernatant from this purification was then removed and additional SPRI beads were added to bring the final overall concentration to 1.0×. DNA bound to these beads was then washed and eluted. The library was quantified using Qubit dsDNA HS Assay Kits (ThermoFisher, # Q32854) and size of the library was verified using Tapestation (Agilent D1000 ScreenTape).

### RT-qPCR for RNA localization analysis

Three wells of the 6-well plate served as one replicate for the RT-qPCR experiments. Cells were fractionated as previously described. 100ng of RNA from each soma and neurite sample were reverse transcribed using LunaScript RT SuperMix (New England Biolabs, #E3010) in a 10µL reaction volume following manufacturer’s instructions. The resulting cDNA was then diluted to 20µL total volume using nuclease-free water. 2µL of diluted cDNA is used as the template for qPCR to estimate the abundances of Firefly and Renilla reporter transcripts or endogenous *Net1* and Tsc1 in the soma and neurite fractions. The qPCR reaction was performed using PrimeTime® Gene Expression Master Mix (Integrated DNA Technologies, #1055772) with differently labeled probe sets for each pair of transcripts, allowing the quantification of two transcripts in the same reaction. Firefly and *Net1* probe sets (Integrated DNA Technologies) are HEX labeled while Rennilla and Tsc1 probe sets (Integrated DNA Technologies) are FAM labeled.

The observed Ct values of the transcripts were within the recommended dynamic range of the assay. Reactions were carried out using a CFX-Opus 384 thermocycler with the following conditions: UNG activation at 50°C for 2 min, followed by polymerase activation at 95°C for 30 s and 40 cycles of 95°C for 5 s, and 60°C for 30 s. Finally, a melting curve was performed by incubating samples at 65°C for 15 s followed by a temperature gradient increase at 0.5°C/s to 95°C. Each sample was measured with three technical replicates. To ensure no contamination, no reverse transcriptase and no template controls were performed. Fold enrichment was calculated using the ΔΔCt method. MIQE guidelines were followed for all qPCR experiments.

### Sequencing analysis

Adapter sequences were removed from reads using cutadapt [58]. Specifically, the sequences GGCGGAAAGATCGCCGTGTAAGTTTGCTTCGATATCCGCATGCTA and CTGATCAGCGGGTTTCACTAGTGCGACCGCAAGAG were trimmed from the 5′ ends of the forward and reverse reads, respectively. A minimum length of 75 was required. The following command was used: `cutadapt -j 16 -g CGGAAAGATCGCCGTGTAAGTTTGCTTCGATATCCGCATGCTA-GGTCGAGGCTGATCAGCGGGTTTCACTAGTGCGACCGCAAGAG --discard-untrimmed --minimum-length 75 -o ${trim1} -p $ {trim2} ${r1} ${r2}`

The trimmed reads were then aligned to the reference oligonucleotide sequences using bowtie2 and the following parameters: bowtie2 -q --end-to-end --no-discordant --no-unal -p 16 -D 150 -x ${ind} −1 ${r1} −2 ${r2} -S $ {sam}

The number of unique UMIs (the first 8 nt of the reverse read) for each reference oligonucleotide was then calculated using https://github.com/TaliaferroLab/OligoPools/blob/master/analyzeresults/UMIsperOligo.py. These UMI counts were then given to DESeq2 [59] to quantify oligonucleotide abundances in each sample and identify sequences enriched in soma or neurite samples.

### Neuronal cell culture, plasmid transfection, and immunostaining

All animal procedures were performed in accordance with the German Welfare for Experimental Animals (LMU Munich, Government of Upper Bavaria). Primary hippocampal neuron cultures were prepared from embryonic day 17 (E17) Sprague-Dawley rat embryos (Charles River Laboratories, Wilmington, MA, USA) as described previously [60]. Briefly, hippocampi were dissected, enzymatically dissociated using trypsin, and mechanically triturated. Dissociated neurons were plated onto poly-L-lysine-coated glass coverslips and maintained in Neurobasal medium supplemented with B27 (Invitrogen, Carlsbad, CA, USA) in a humidified incubator at 37 °C and 5% CO₂.

The plasmids pUBC-NLS-HA-stdMCP-stdGFP and pRSV-LacZ-128xMS2 have been described previously [34,61]. To generate the MS2 reporter constructs used in this study, 260-nt sequences corresponding to *Net1* WT, *Net1* mutant, *Trak2* WT, and *Trak2* mutant were cloned into the BamHI restriction site in the 3′ untranslated region (3′UTR) of the pRSV-LacZ-128xMS2 reporter plasmid. Neurons were transiently transfected at 15-16 days in vitro (DIV) using calcium phosphate co-precipitation (1 µg pUBC-NLS-HA-stdMCP-stdGFP + 2 µg of each MS2 reporter plasmid) and fixed after 16 hours of expression for 10 min with 4% PFA [60]. Neurons were immunostained with a primary mouse anti-Tau antibody (Synaptic Systems; 1:2000), followed by a donkey anti-mouse Alexa Fluor 555-conjugated secondary antibody (Life Technologies; 1:1000), to facilitate the distinction of axons and dendrites.

### Imaging and data analysis of primary neurons

Images were acquired using a Zeiss Axio Observer Z1 inverted microscope equipped with a 63× Plan-Apochromat oil-immersion objective (NA 1.40), a Colibri LED light source, and an Axiocam 506 mono camera, controlled by Zen software (Zeiss, Oberkochen, Germany). For analysis of the MS2 reporter assay, dendrites or axons were selected and straightened using the segmented line tool in ImageJ. Individual MS2 particles were manually identified using the multipoint tool. Distances from the soma and particle counts were quantified using a custom script that extracted the position of each particle along the neuronal process.

### 1M7 acylation of RNA localization elements for structure probing

For each purified RNA, 10 pmol was diluted into 14 µl RNase-free water, denatured at 95 °C for 2 minutes, and cooled on ice for 2 minutes. Denatured RNA was combined with 4 µl of 5× folding buffer to achieve a final 1.1× concentration [110 mM HEPES pH 8.0, 110 mM NaCl, 5.5 mM MgCl_2_] and was folded to equilibrium for 20 min at 37 °C on a heat block. During incubation, a fresh stock of 100 mM 1M7 (1-Methyl-7-nitroisatoic anhydride, 73043-80-8, MilliporeSigma) was prepared by dissolving powder in neat anhydrous DMSO. Two new empty tubes were prepared for each 1M7 and mock treatment: one tube with 1 µl of 100 mM 1M7, the other with 1 µl of DMSO. After incubating RNA, treatment tubes were placed on the 37 °C heat block and 9 µl of folded RNA was added to 1M7 and DMSO treatment tubes (adequate mixing upon dilution), resulting in 1× folding buffer reactions [100 mM HEPES pH 8.0, 100 mM NaCl, 5 mM MgCl_2_] with 10 mM 1M7 or 1.4 mM DMSO, respectively. The samples were incubated for 2 min (in multiple excess of the 17 s aqueous half-life of 1M7) at 37 °C. The RNA samples were exchanged into Tris-EDTA buffer over G-50 sephadex columns (illustra™ MicroSpin™, Cytiva), 2 minutes at 735 × g according to manufacturer’s protocol.

### MaP RT (mutational profiling reverse transcription)

In 10 µl of total volume, 2-3 pmol of RNA (7 µl of G-50 purified RNA product) was denatured at 70 °C for 5 min in a mixture containing 20 nmol of each dNTP base and 2 pmol of RT primer complementary to the 3’ end of the RNA constructs. After 5 min, the mixture was cooled to 4 °C for 2 min. 9 µl of freshly made 2.22× MaP buffer (111 mM Tris-HCl (pH 8.0), 167 mM KCl, 13.3 mM MnCl_2_, 22 mM DTT and 2.22 M betaine) was then added to the mixture, which was then equilibrated at 25 °C for 2 min. Subsequently, 1 µl (200 units) of SSIIRT (SuperScript™ II Reverse Transcriptase, Invitrogen) was added to the reaction, which was then placed in a thermocycler with the following settings: 10 min at 25 °C, 90 min at 42 °C, 10 cycles of 2 min at 42 °C and 2 min at 50 °C, and finally 10 min at 70 °C to heat inactivate the SSIIRT.

### SHAPE-MaP library preparation and sequencing

Targeted amplicon sequencing libraries were generated by two-step PCR. First, 1 µl of cDNA from the MaP RT reaction was used as a template for PCR with 1.25 µL each of 10 µM construct-specific forward and reverse primers (with extended Illumina-specific sequences) and Q5 hot-start polymerase (NEB) in a total reaction volume of 25 µL. The reactions were performed in a thermocycler with the following conditions: 1 min at 98 °C, 15 × (10 s at 98 °C, 30 s at 64 °C, and 30 s at 72 °C), and 2 min at 72 °C. PCR products were purified by capture with 0.8× volumes of SPRI beads (Mag-Bind® TotalPure NGS, Omega Bio-Tek), beads were washed twice with 80 % ethanol, and libraries were eluted with 15 µl of nuclease-free water. 2 ng of step 1 PCR products were used as templates for step 2 PCR with 5 µL each of 5 µM i5/i7 indexing primers and Q5 hot-start polymerase (NEB) in a total reaction volume of 50 µL. The reactions were performed in a thermocycler with the following conditions: 1 min at 98 °C, 10 × (10 s at 98 °C, 30 s at 67 °C and 30 s at 72 °C), and 2 min at 72 °C. PCR products were purified with 0.65× volumes of SPRI beads, beads were washed twice with 80 % ethanol, and libraries were eluted with 15 µl of nuclease-free water. Size distributions and purities of amplicon libraries were verified (4150 TapeStation, Agilent). About 120 amol of each library was sequenced on a MiSeq instrument (Illumina) with 2 × 250 paired-end sequencing.

### Analysis with Shapemapper2 and RNAStructure

FASTQ files from sequencing runs were input into ShapeMapper2 (defaults) for read alignment to each localization element sequence and mutation counting, including --amplicon and --primers flags to exclude internal reverse transcription initiation sites and excluded primer regions from analysis. Normalized reactivity values from ‘profile.txt’ outputs generated by Shapemapper2 were used for downstream structural modeling. Minimum free energy models of RNA secondary structures were generated from normalized SHAPE reactivities using RNAstructure Fold with default folding parameters and the -mfe and -sh flags. Pairing probabilities were generated by applying a partition function using a combination of RNAStructure functions “partition” and “ProbabilityPlot” (with -sh flag)[62]. SHAPE reactivities and pairing probabilities were plotted on each construct sequence using arcPlot (https://github.com/Weeks-UNC/arcPlot).

### Comparison of wildtype and mutant localization element SHAPE reactivities

SHAPE modification rates of wildtype and mutant localization element RNAs were normalized to each other using a median difference minimization strategy [63,64] to enhance sensitivity to single-nucleotide level differences. First, the log relative reactivities for each dataset were calculated as follows:

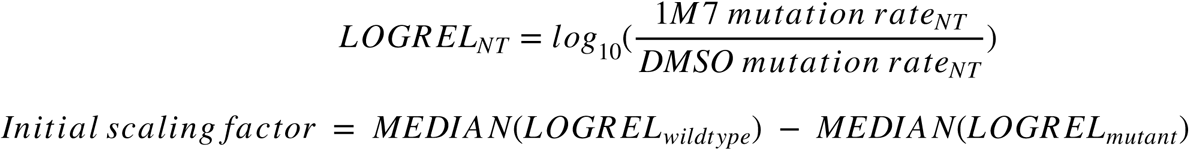

The LOGREL_mutant_ values were adjusted up by the initial scaling factor, and differences were calculated for each nucleotide:

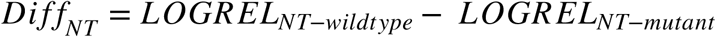

The final scaling factor (added to mutant LOGREL values) was calculated as the value that minimizes the median for all nucleotides of |Diff_NT_|. New Diff_NT_ values were computed with the final scaling factor, and Z-scores were computed for each nucleotide:

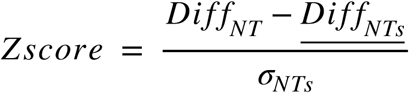

Only nucleotides with Z-scores > 2.576 standard deviations from the mean (99% confidence intervals) were considered significant shifts in SHAPE reactivity.

### MaP primers

localization element RT primer (reverse) 5′-CACTAGTGCGACCGC

localization element i7 MaP step 1 primer (forward)

5′-GTCTCGTGGGCTCGGAGATGTGTATAAGAGACAGNNNNNGCTTCGATATCCGCATGC

localization element i5 MaP step 1 primer (reverse)

5′-TCGTCGGCAGCGTCAGATGTGTATAAGAGACAGNNNNNCACTAGTGCGACCGC

i7 index step 2 primer

CAAGCAGAAGACGGCATACGAGAT-index-GTCTCGTGGGCTCGG

I5 index step 2 primer

AATGATACGGCGACCACCGAGATCTACAC-index-TCGTCGGCAGCGTC

## AUTHOR CONTRIBUTIONS

Experiments were conceived by CM, CW, AWM, DD, MK, and JMT and were performed by CM, CW, SSS, KB, and AM. Funding for the project was secured by CW, AWM, DD, MK, and JMT. The manuscript was written by CM and JMT and edited by CM, CW, MK, and JMT.

## RESOURCE AVAILABILITY

### Lead contact

Further information and requests for resources and reagents should be directed to and will be fulfilled by the lead contact, J. Matthew Taliaferro.

### Materials availability

All materials generated in this study are available through the lead contact upon request.

### Data availability

All high-throughput sequencing data associated with these experiments has been deposited at the Gene Expression Omnibus under accession number GSE334718.

## SUPPLEMENTARY TABLES

**Table S1**. Neurite enrichments for all oligonucleotides in the sufficiency MPRA in the GFP and firefly luciferase reporter contexts.

**Table S2**. Neurite enrichments for all oligonucleotides in the deletion MPRA in the GFP and firefly luciferase reporter contexts.

**Table S3**. Neurite enrichments for all oligonucleotides in the mutation MPRA in the GFP and firefly luciferase reporter contexts.

**Table S4**. Neurite enrichments for all oligonucleotides in the shuffle MPRA in the GFP and firefly luciferase reporter contexts.

**Table S5**. Neurite enrichments for all oligonucleotides in the SHAPE MPRA in the GFP and firefly luciferase reporter contexts.

## SUPPLEMENTARY FILES

**Supplementary File 1**. Fasta file of all oligonucleotide sequences used across all MPRAs. The name of each oligo contains an abbreviation indicating what MPRA it is from following a “+”. For the sufficiency MPRA, the abbreviation is “suff”, for necessity “ness”, for mutation “mut”, for shuffle “shuff”, and for SHAPE “shape.” Within the sufficiency MPRA, the oligo names indicate the parent LE sequence an oligo is derived from, and the start and end indices of the subset of the parent LE being tested as follows: parentLEname_start|end. These indices are inclusive.

For the necessity MRPA, the oligo are named as follows: parentLEname_start:end, with start and end indicating the start and end of the indices deleted from the parent LE sequence. If the same deletion window led to the same oligo sequence, the name of that sequence were the names of the oligos with windows that led to the same sequence separated by “$,” i.e. parentLEname_start:end$parentLEname_start:end. As in the sufficiency MPRA, these indices are inclusive.

For the mutation MPRA, most oligos are named: parentLEname_start:end_rep. Start and end are the inclusive start and end indices of the mutation window, and rep is used to indicate the set of mutations to a single window. Additionally, mutations were made to the *Trak2*, *Net1*, and *Trp53inp2* LEs based on their minimum free energy structures. Compensatory mutations made based on these structures are named: parentLEname_startIindex:endIindex::startJindex:endJindex_rep. Mutations were also made to the sequences flanking the CEs of these oligos. These were named parentLEname_flanks-rep.

For the shuffle MPRA, oligos are named: parentLEname_start:end_rep_h-hammingDistance. Shuffles were also made to the sequences flanking the CEs of these LE. These were named: parentLEname_flanks-rep.

For the SHAPE MPRA, oligos are named: parentLEname_stem-helixGroupNumbersSeparatedBy“-”mutationType_rep. Where the mutation types are i, j, or comp (compensatory). Note that in figures the i and j mutation types are swapped to follow naming conventions of having the more 3′ side of a helix the i side, and the more 5′ the j. Mutations were also made to the terminal loops and bulges in the SHAPE-informed structures, these were named: parentLEname_loop-loopNumberSepBy“-”repN and parentLEname_bulge-BulgeNumberSepBy“-”repN.

## SUPPLEMENTARY FIGURES

**Figure S1.**
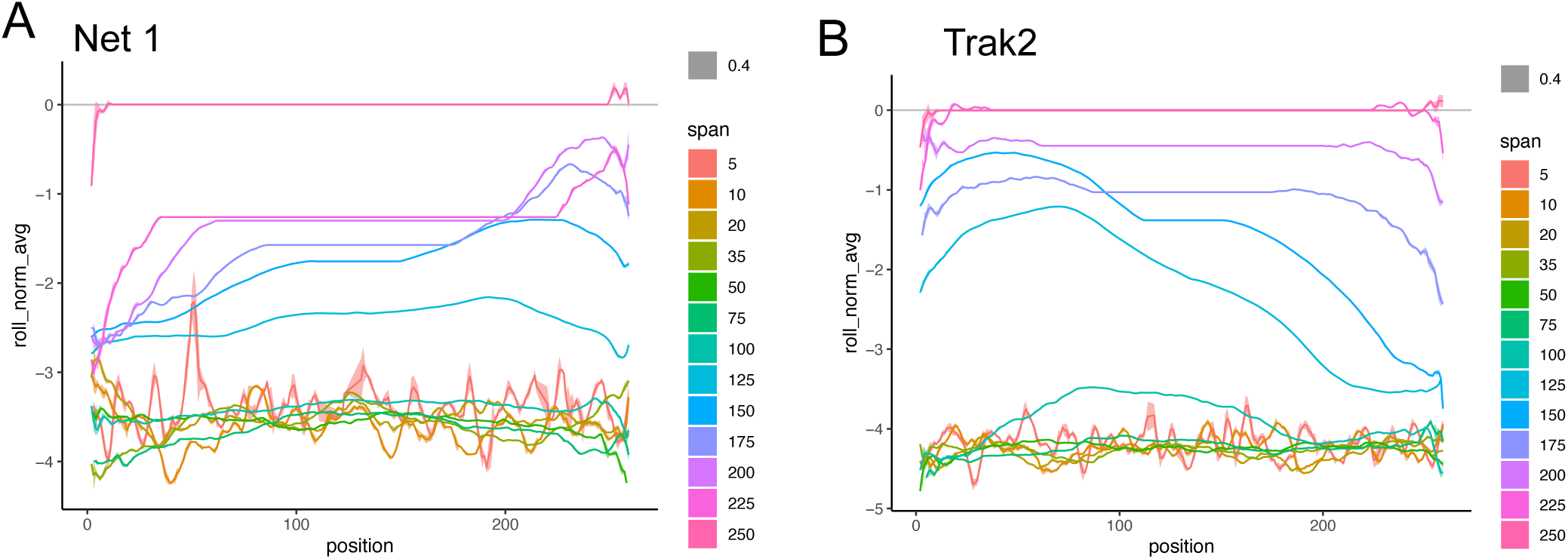
(A) Neurite enrichment for reporters of all lengths tested of *Net1* LE sequence subsets derived from the parent 260mer LE. Rolling mean, k=3. Ribbon represents standard error. (B) As in A, but for *Trak2* LE.

**Figure S2.**
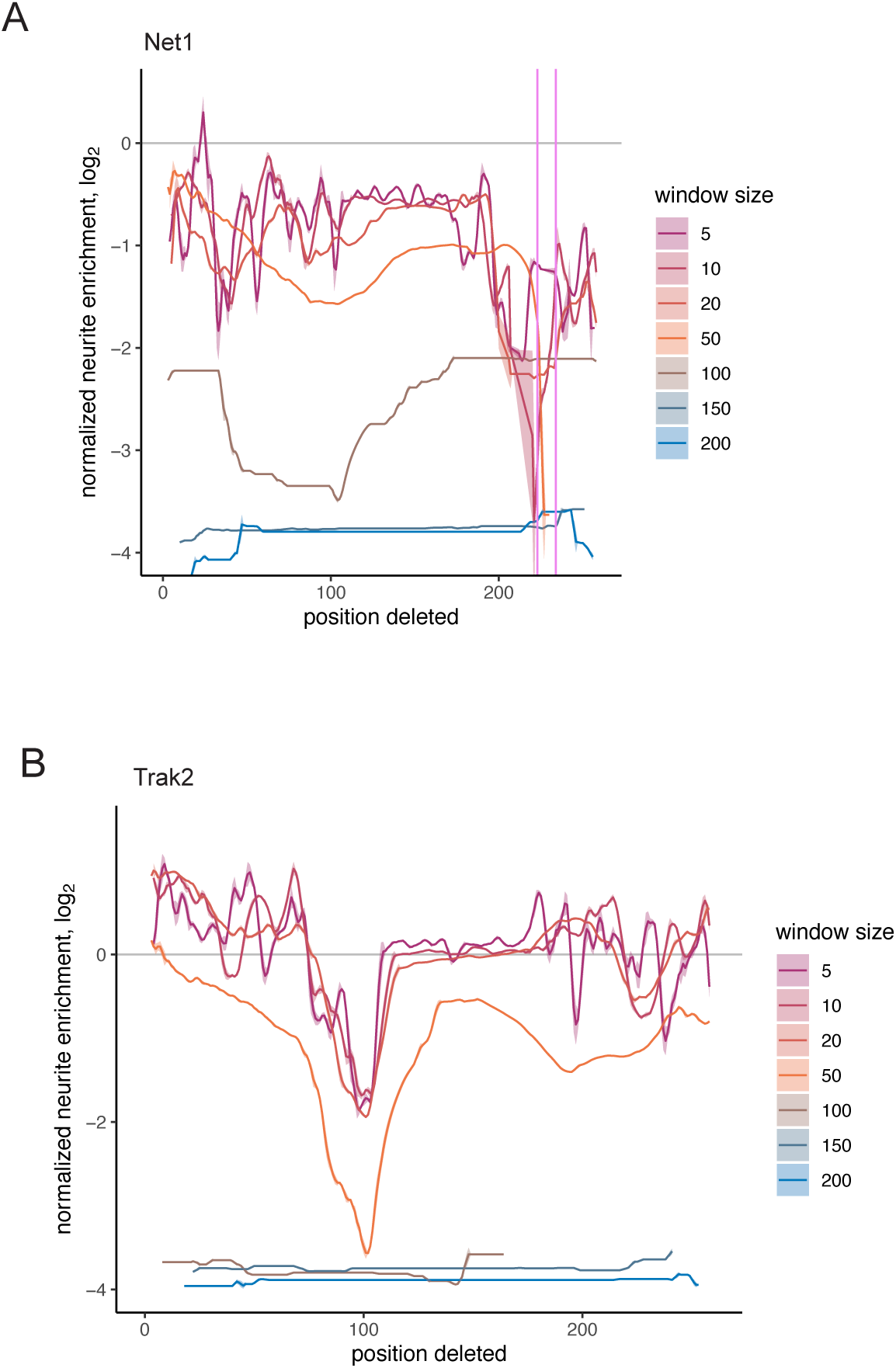
(A) Neurite enrichment for reporters containing *Net1* LE derived sequences containing all tested deletion window sizes. Rolling mean, k=3. Ribbon represents standard error. (B) As in A, but for *Trak2* LE.

**Figure S3.**
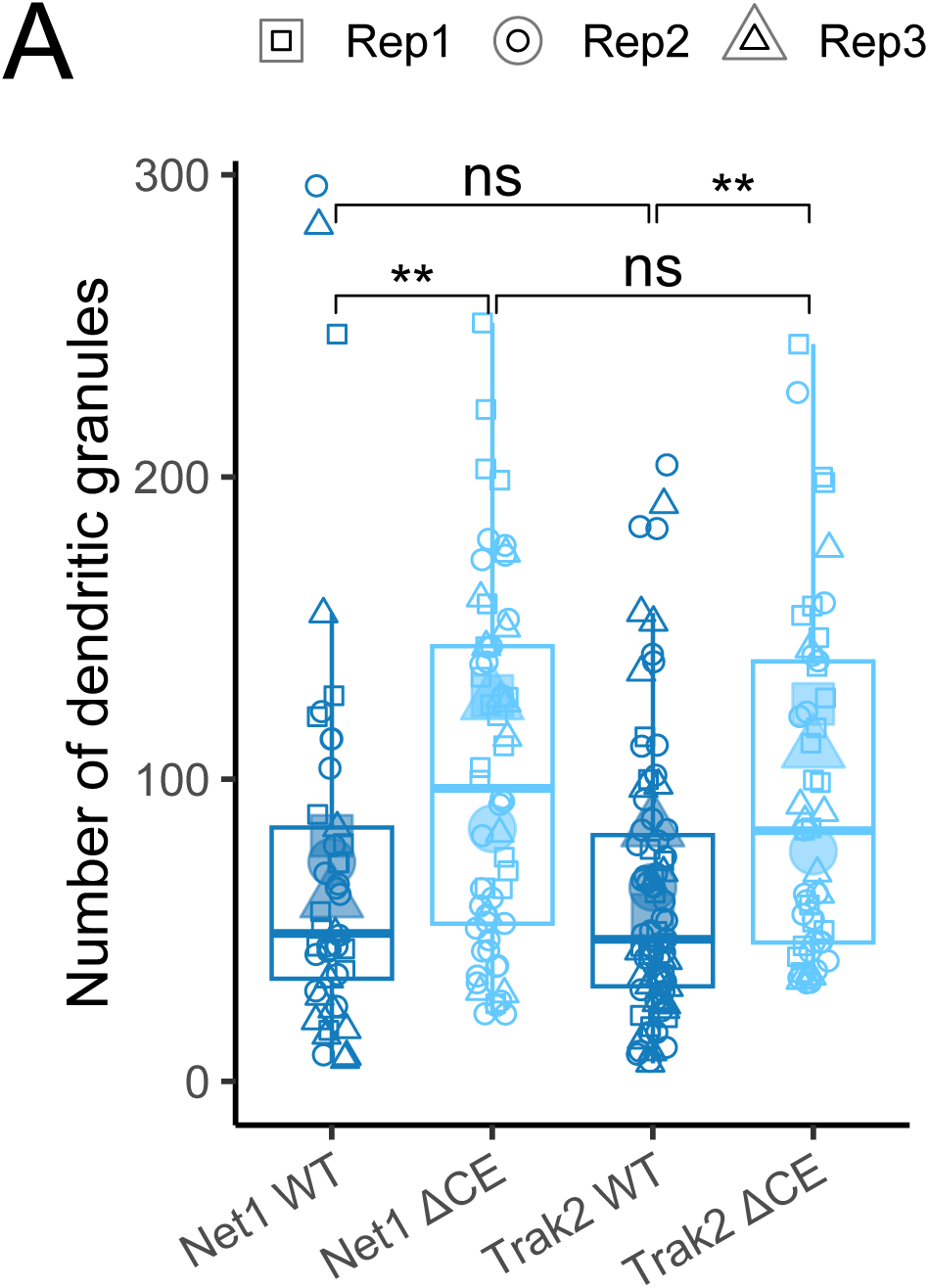
(A) Quantification of the number of LE containing-granules in the dendrites of rat hippocampal neurons.

**Figure S4.**
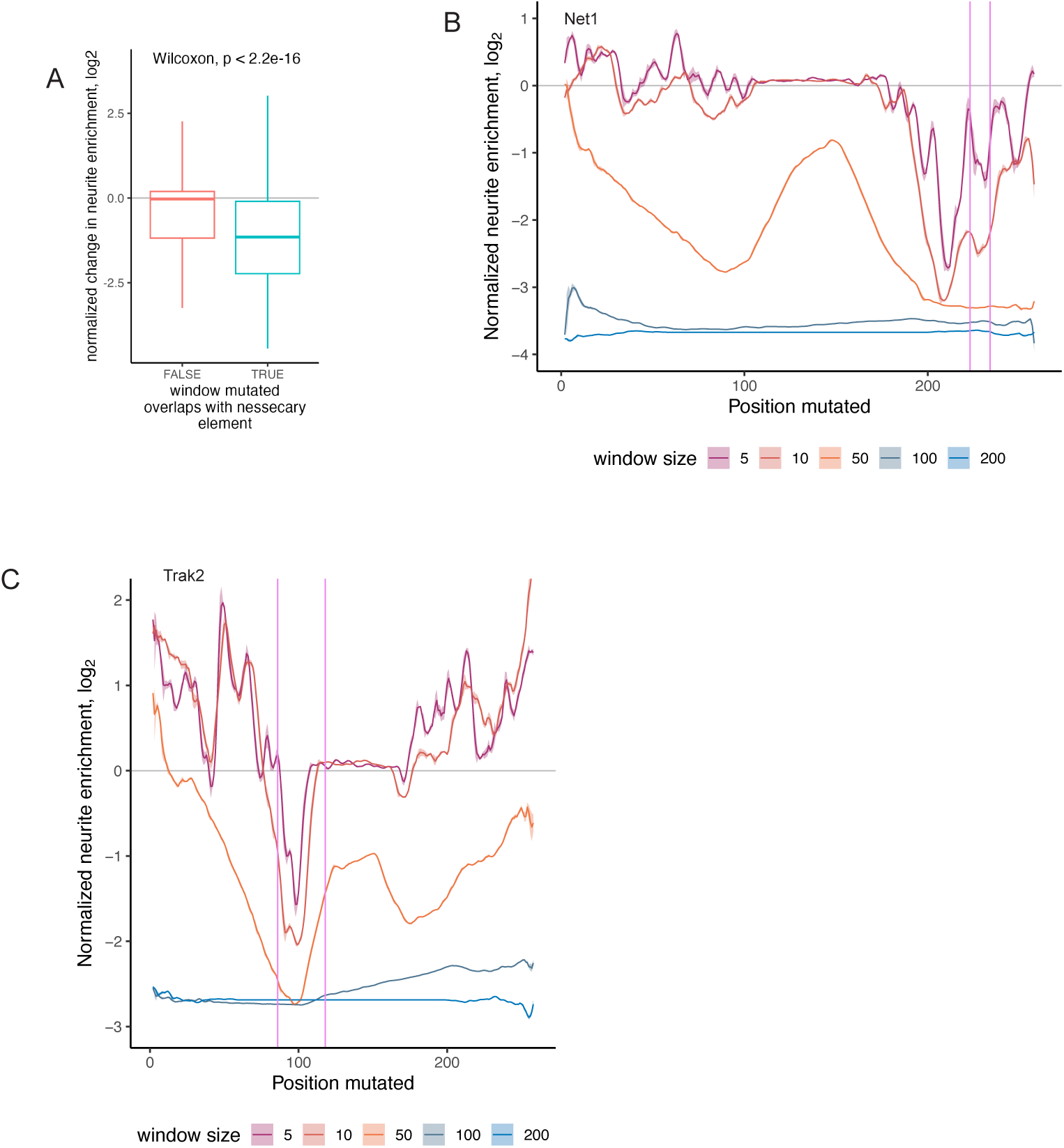
(A) Differences in neurite localization of oligos that do or do not have mutations within their CE. All deletion window sizes were pooled. Oligos were derived from *Net1*, *Trak2*, and *Trp53inp2*. (B) Neurite enrichment for reporters containing *Net1* LE derived sequences containing all tested mutation window sizes. Rolling mean, k=3. Ribbon represents standard error. (C) As in B, but for *Trak2* LE.

**Figure S5.**
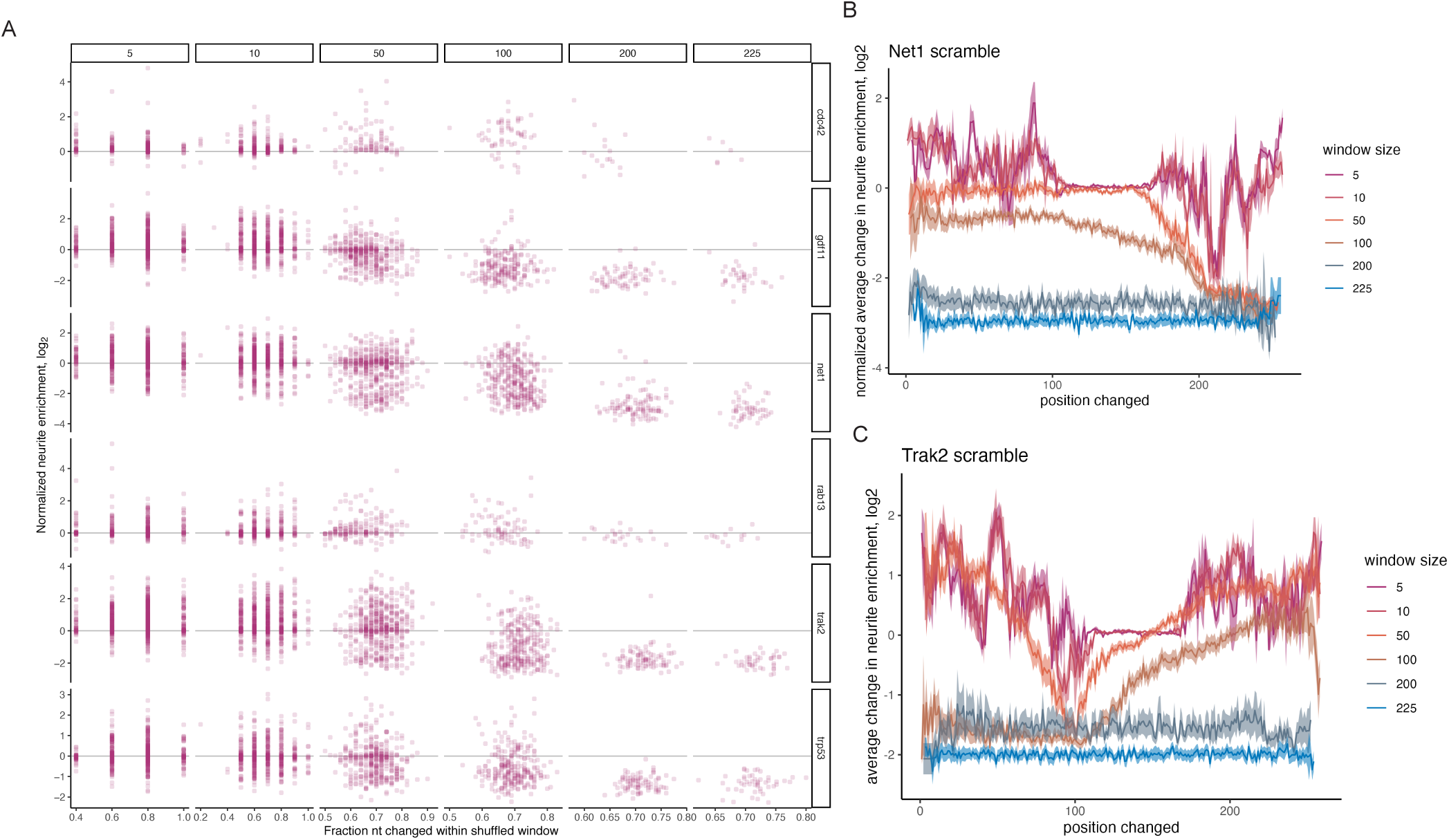
(A) Neurite enrichment of reporters from all LEs depending on the size of the window shuffled and the percentage of nucleotide identities within the changed. Plot is faceted such that window size is represented by columns and parent LE is represented by rows. The full gene names of the parent genes are, from top to bottom: *Cdc42bpg*, *Gdf11*, *Net1*, *Rab13*, *Trak2*, and *Trp53inp2*. (B) Neurite enrichment for reporters containing *Net1* LE derived sequences containing all tested shuffle window sizes. Rolling mean, k=3. Ribbon represents standard error. (C) As in B, but for *Trak2* LE.

**Figure S6.**
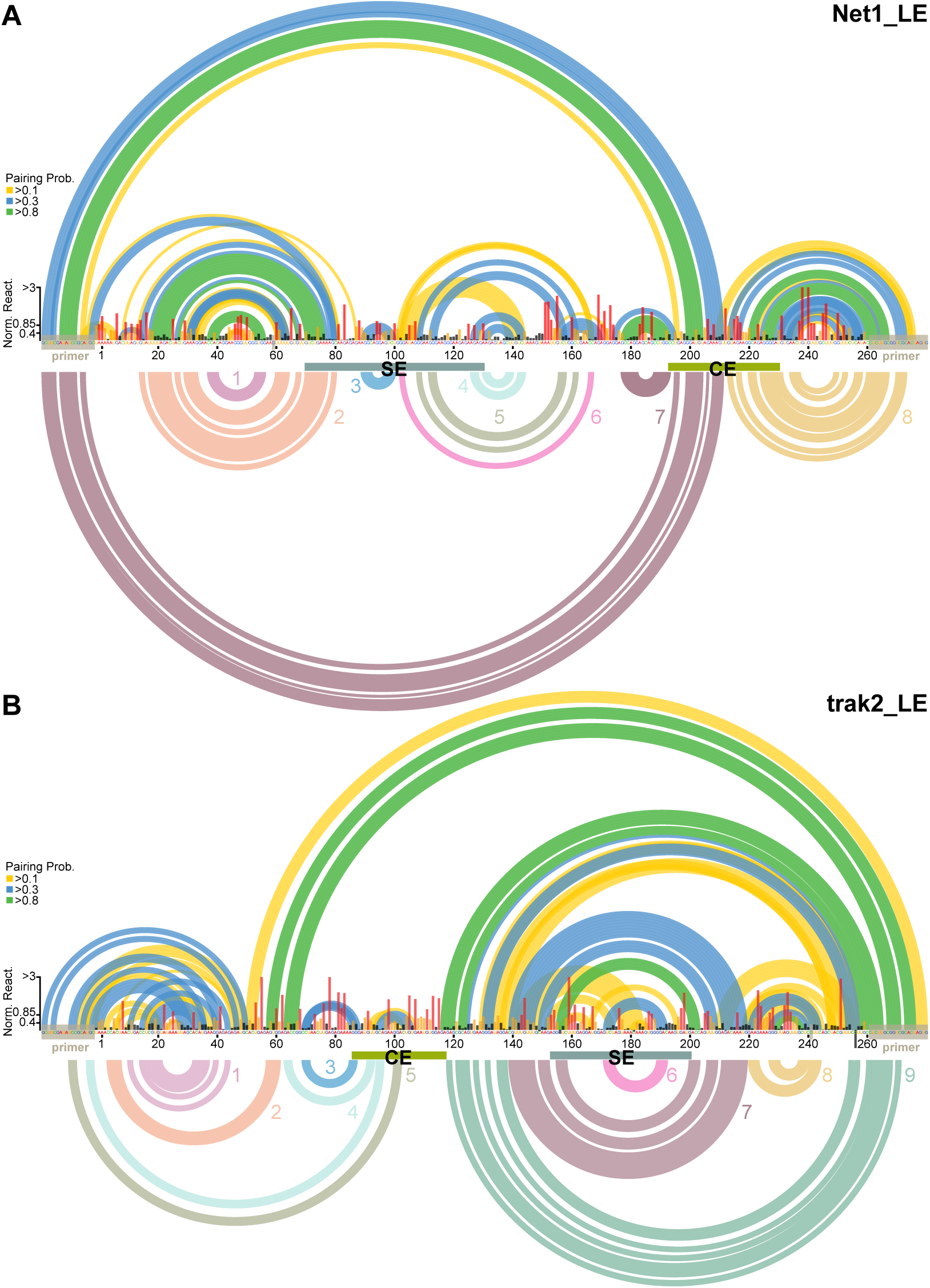
SHAPE-informed free energy modeling for the *Net1* (A) and *Trak2* (B) localization elements. Support element (SE) and critical element (CE) regions are labeled, and nucleotide position and base identity are displayed. Normalized SHAPE reactivity for each nucleotide is plotted as columns (black < 0.4 < yellow < 0.85 < red), and primer regions and high background sites that do not contribute SHAPE data for modeling are labeled with gray bars. Pairing probabilities derived from partition function are shown as colored arcs above sequences according to indicated thresholds (pairing probabilities less than 0.1 are not displayed). Minimum free energy model base-pairing explored by mutation in Figure 6 are labeled and displayed in corresponding colors below sequences.

